# DIRTNet: Enabling Root Phenotyping with Fiber Bragg Grating Sensors

**DOI:** 10.64898/2026.09.22.753535

**Authors:** Kabir Hossain, Steven Binder, Mable Fok, Alexander Bucksch

## Abstract

Non-destructive methods for real-time phenotyping of root growth and development are essential to accelerate the breeding of climate-resilient crops. We present DIRTNet (Digital Investigations of Root Traits Net), a hybrid deep learning model that interprets in-soil signals collected by Fiber Bragg Grating (FBG) sensors. We first evaluated DIRTNet in a controlled simulation study in which metal rods of varying diameters were inserted into soil to emulate root growth. Using augmented training data with an 80:20 train–test split, DIRTNet surpassed baseline models, achieving 0.97 accuracy for root depth prediction and 0.94 for root diameter prediction. We then applied the best-performing models to maize (Zea mays) roots monitored by six FBG sensors placed at different soil location and depths. A control trial and a drought-stress trial yielded 192 samples, split 80:20 into training and holdout sets; the hold-out portion was divided equally into validation and test sets. Augmentation was applied only to the training data. On real maize root data, DIRTNet achieved 0.92 accuracy for both root depth and root width prediction across variable growth periods, covering root mass lengths up to 20 cm and diameters up to 10 cm. The prediction accuracy is independent of the soil moisture and drought condition. In a fixed-interval analysis, accuracy reached 0.97 for days 25–42 after planting and 0.93 for days 25–48, demonstrating robust performance as root complexity increased. DIRTNet also classified control versus drought-stress conditions with 0.87 accuracy. Together, these results establish DIRTNet as a continuous, scalable, and non-destructive approach for root phenotyping with strong potential to support crop improvement under changing environmental conditions.

## 1. Introduction

Rising temperatures and increasingly frequent extreme weather events are accelerating soil degradation and reducing the land available for cultivation. Up to 40 percent of the world’s land area is already degraded, affecting nearly half of humanity [1]. These mounting edaphic stresses, compounded by unsustainable farming practices [2], pose a serious threat to agricultural productivity [3]. Environmentally, soil degradation significantly affects soil organic carbon levels, which are crucial for soil fertility and agricultural productivity [4, 5]. As a result, crop production is expected to become more challenging, and innovative solutions are needed to ensure food security and sustainable agricultural practices. One of the critical factors contributes the nutrient and water uptake efficiency of plants [6, 7] is the Root System Architecture (RSA), which determines how well a plant performs in different stress environments [8]. Therefore, the phenotyping of RSA can enable valuable solutions for addressing today’s global challenges, such as enhancing crop resilience to improve food, feed and fiber security. Addressing these challenges requires highly interdisciplinary approaches, including the development of new technologies that can quantify the formation of root phenotypes in relation to their growth environment [9]. Unfortunately, these technologies remain largely inaccessible because of their high cost, technical complexity, and limited availability.

Traditional methods for studying RSA, such as MRI [10], X-ray computed tomography [11, 12, 13], and Ground-Penetrating Radar (GPR) [14], although effective, are often expensive, resolution limited to capture the critical phase of seedling establishment, and not suitable for continuous monitoring over several months of root development. Additionally, methods relying on manual excavation with Shovelomics [15, 16, 17, 18], are labor-intensive and destructive to the plant, limiting their utility for large-scale continuous measurements of RSA development. There is an urgent need for nondestructive, cost-effective, and continuous root monitoring technologies to gain insights into root development.

Fiber optic sensors are compact in size, robust to harsh environments, can be remotely operated and continuously take measurements in time. Some fiber optic sensors can even provide continuous measurement in space using a single piece of sensing fiber [19]. Various science disciplines already employ fiber optic sensors [20], including structural health monitoring, biomedical applications, earth sciences, and aerospace applications. Fiber optic sensors stand out due to their high sensitivity in detecting small changes, compactness, immune to electromagnetic interference, and multiplexing ability [20]. As such, fiber optics has great potential as the enabling technology towards accurate, non-invasive measurement of root growth dynamics, offering continuous insights into root development across different environmental conditions.

Recently, Tei et al. [21] presented a method using distributed fiber optics sensors for non-destructive, real-time root monitoring in artificial growth media. They demonstrated the detection of sub-millimeter diameter object penetration in artificial growth media. In their experiment, artificial roots were used to simulate the penetration of real plant roots to characterize the fiber sensor response. They also developed a computational model to visualize the roots of tuber crops and monocotyledons. Their system, however, requires a relatively complex sensing setup comprising an optical fiber attached to a perforated polymer film, a spiral supporting structure, and a commercial optical reflectometer. Moreover, the experiments were conducted in a soil-like substrate rather than natural soil, which may not adequately represent the heterogeneity and complex root–soil interactions found under field conditions.

Our prior work [22, 23, 24] introduced a proof-of-concept in-soil FBG sensing system for root phenotyping using pseudo-root experiments and a preliminary single-plant maize study under controlled conditions. This study established the feasibility of in-soil FBG sensing for non-destructive root trait estimation and highlighted its potential for continuous root monitoring. Nevertheless, traditional neural networks, such as Deep Convolutional Network (CNN) [25], Visual Geometry Group (VGG) [26], and Residual Networks (ResNet) [27] are not designed to address the combined temporal and spatial information processing needed for continuous root phenotyping with fiber optics sensors. Moreover, fiber optic sensing signals are often weak, noisy, and indirectly related to root growth dynamics, further limiting the effectiveness of standard feature extraction approaches [21]. Therefore, we developed a hybrid deep learning model for root phenotyping named DIRTNet. The newly developed DIRTNet combines the strength of Residual Networks (ResNet) and Visual Geometry Group (VGG) components for feature extraction with the capabilities of Gated Recurrent Unit (GRU) components to capture temporal dependencies [28].

In our experiments, we first simulated root growth with pseudo-roots by inserting metal rods of varying diameters into the soil, enabling the controlled generation of root-like strain signals measured by fiber Bragg grating (FBG) sensors. These simulation data allowed us to systematically benchmark DIRTNet against ten baseline models for predicting root depth and diameter. The results demonstrated that DIRTNet consistently achieved the highest accuracy, precision, and recall, along with the lowest RMSE among all compared models. These controlled experiments established a strong foundation for validating the effectiveness of the proposed approach.

Building on the simulation results, we conducted a second experiment that uses FBG sensors to monitor actual root growth in maize, under both control and drought conditions. The collected sensing data were used to train and evaluate DIRTNet for trait estimation of real root and classification of environmental conditions. By continuously analyzing FBG signals, DIRTNet captures drought induced changes in root growth patterns, supporting both root trait prediction and the classification of growth conditions. This experiment demonstrates the ability of DIRTNet for agriculture and enables direct evaluation of root responses to drought stress.

## 2. Methodology

### 2.1. Growing conditions

The same maize cultivar (Early Golden Bantam) was used in all real root growth experiments. Each plant was grown in a 5-gallon bucket containing about 5.9 kg of dry potting soil.

Two experimental trials were conducted. The first trial included five plants grown under well-watered condition. The second trial included six plants, of which three were maintained under well-watered condition and three were subjected to drought stress.

In well-watered condition, the plants were irrigated with 150 mL of water every 3 days to maintain consistent soil moisture throughout the 75-day growth period. For drought-treated plants, the same irrigation schedule was applied for the first 25 days, after which no further water was provided for the remaining 50 days.

On day 75^th^, all plants were carefully removed from the soil, and root system traits, including total root depth and root width, were manually measured for ground-truth validation.

### 2.2. FBG Principle and Experimental Setup

A FBG optical sensor is a piece of optical fiber with a short region where core refractive index is changed periodically [29], forming a grating with a specific Bragg wavelength. When broadband light enters the FBG, a specific Bragg wavelength is constructively interfered due to the grating period and is reflected back to the FBG input while the rest of the wavelength will pass through the FBG. Strain applied on the FBG would change the grating period and results in a change of Bragg wavelength. Therefore, a shift in Bragg wavelength can be observed when root expansion occurs or when soil particles shrink due to drought conditions. The shift in FBG’s Bragg wavelength can be monitored by aligning a single wavelength laser’s wavelength to match the Bragg wavelength, such that the induced wavelength shift is directly translated to a change in reflected optical power.

#### 2.2.1. Pseudo-root experimental setup

The detail description of the pseudo-root set up for root depth and diameter measurement can be found in our previous work [23]. The optical sensing setup consists of three FBGs with Bragg wavelengths of 1550.0nm, 1552.5nm, and 1553.3nm are buried within soil near the pseudo-root insertion zone. Each FBG is connected to their own single wavelength laser diode (LD), optical circulator, and optical power meter (PM). At the beginning of each experiment, each laser diode is tuned to a wavelength corresponding to a power measurement 3 dB lower than the maximum possible reflected power. Ensuring that we operate in the most linear region of the FBG reflection spectrum. Inserting the pseudo-root induces strain on each FBG sensor and the resulting optical power per sensor is recorded with the software NI LabView in a text file for later analysis with DIRTNet.

#### 2.2.2. Real maize experimental setup

We conducted two experiments monitoring real root growth of potted maize plants.The first trial consisted of five plants grown under control cond ition with two layers of FBGs embedded below the seed. The first layer was 5 cm below the seed with one FBG directly centered beneath and two FBGs 2 cm’s to the left and right. The second row was 20 cm below the seed and was also comprised of the same left, center, and right layout. The second trial was comprised of 6 plants with 3 subjected to drought condition and 3 subjected to control condition. The FBG placement was similar to the first trial, however the left and right FBGs were 10 cm away from the center. This wider spacing helped capture broader root-growth and drought-related strain variations, providing more spatial information for DIRTNet training. The detailed configuration of the real maize root setup is shown in Figure 1.

**Figure 1:**
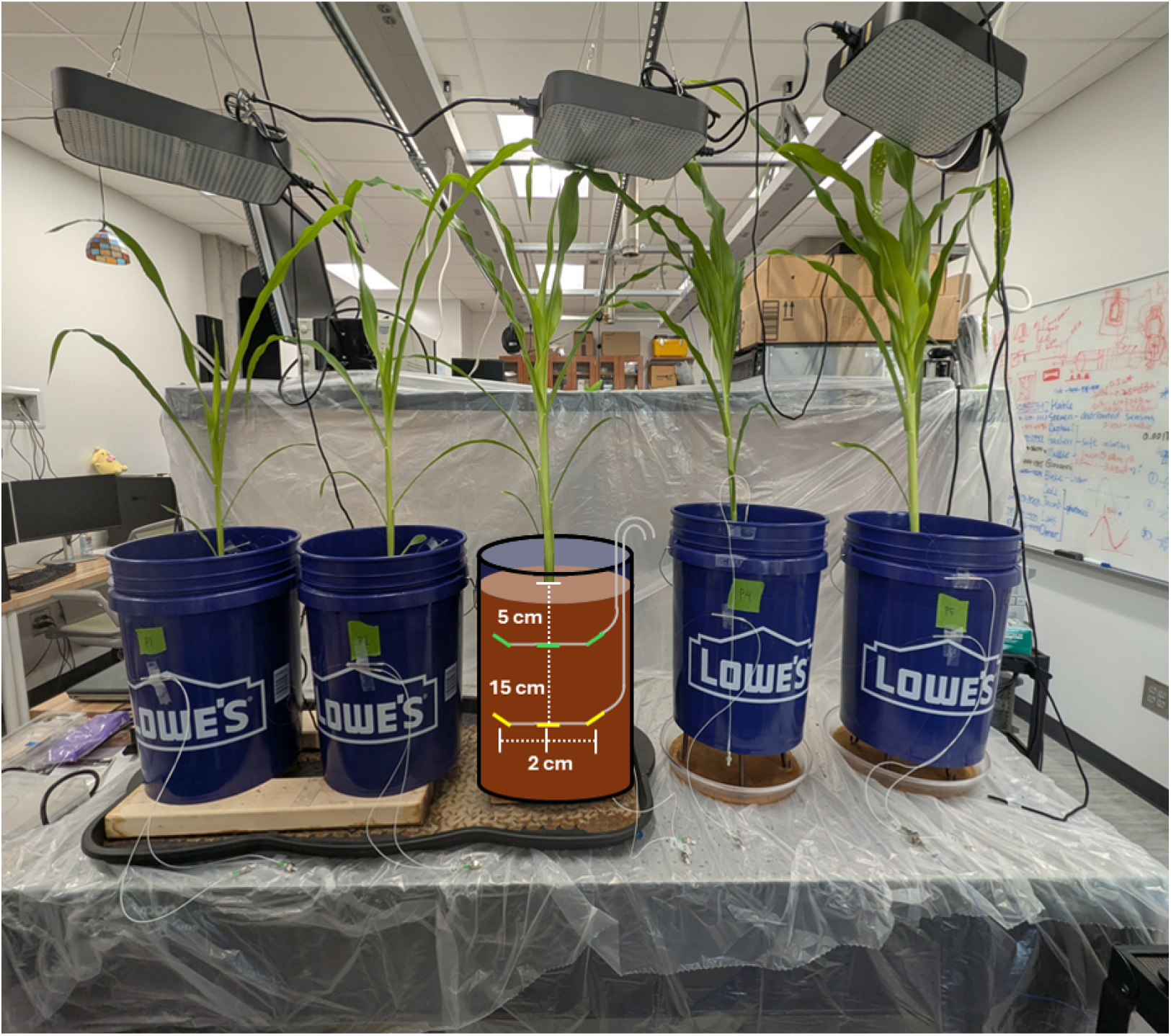
Illustration of the maize root monitoring system with embedded FBG sensors for non-destructive phenotyping.

Every day after planting, the reflection spectrum of all FBGs was recorded using a broadband light sourse and an optical spectrum analyzer. This was done to compensate the influence laser frequency drift may have on the measurement. Using the first acquisition day’s optical spectrum, the wavelength corresponds to each FBGs 3 dB point on the shorter reflected wavelength side was extracted. Then the power measured at that wavelength across all measurement days was extracted for use in subsequent analysis and to simulate a reflected optical power measurement system.

### 2.3. DIRTNet Development

DIRTNet consists of five components (Figure 2). Leverages architecture elements of ResNet and VGG are for addressing spatial feature extraction and architecture elements of GRU is for accounting for temporal features in the FBG data.

**Figure 2:**
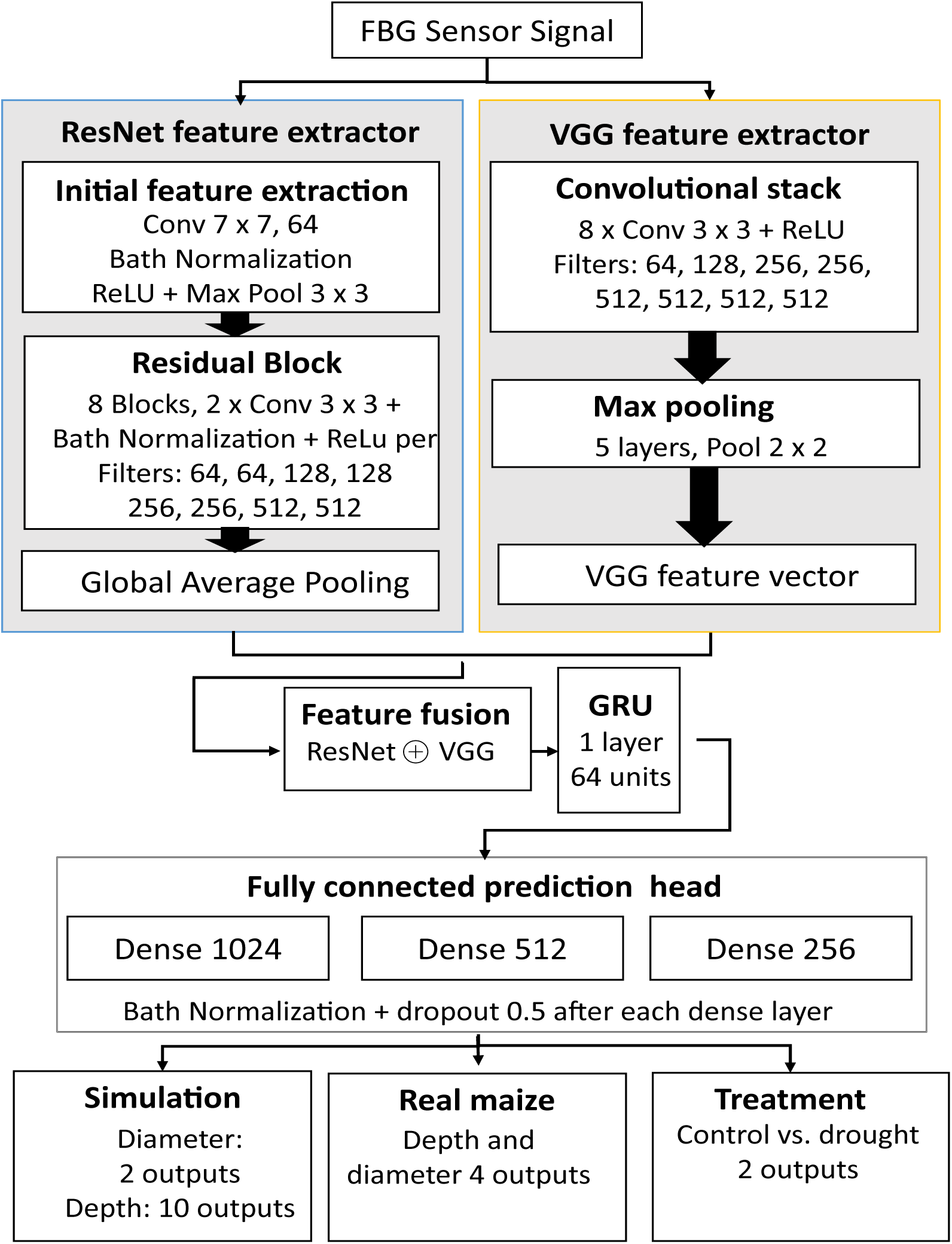
Architecture of the proposed DIRTNet consisting of five main components: ResNet feature extractor, VGG feature extractor, feature concatenation, GRU temporal modeling, and fully connected prediction layers.

The first component of DIRTNet is the part of the ResNet architecture and designed to extract high-level spatial features from the FBG signal. This component consists of an initial convolution layer with a 7×7 kernel size and 64 filters, followed by one batch normalization layer, one ReLU activation layer, and one max pooling layer with a 3×3 pool size. Subsequently, eight residual blocks are employed. Each residual block contains two convolution layers with 3×3 kernel size and filter sizes of 64, 64, 128, 128, 256, 256, 512, and 512. Each convolution layer is followed by one batch normalization layer and one Rectified Linear Unit (ReLU) activation. Finally, a global average pooling layer is applied.

The VGG component contributes additional spatial feature extraction. It includes eight convolution layers with a 3×3 kernel size and filter sizes of 64, 128, 256, 256, 512, 512, 512, and 512, with additional five max-pooling layers with a 2×2 pool size and eight ReLU activation layers.

From an architectural perspective, the ResNet branch integrates information over large receptive fields through deep residual learning, emphasizing global and low-frequency spatial patterns in the FBG signals, such as overall signal magnitude trends, spatial variance across sensors, and smoothly varying strain patterns distributed across multiple sensing locations. In parallel, a VGG-based convolutional branch is used to extract spatial features through a sequence of convolutional layers with small kernel sizes. The VGG architecture provides a complementary spatial information by focusing on local spatial correlations through small convolutional kernels and sequential layer stacking. This design emphasizes short-range, higher-frequency spatial variations in the FBG signals, such as local spatial gradients, short-range fluctuations, and localized changes in strain measurements. Although ResNet and VGG operate on the same inputs, their different depths and receptive-field growth introduce distinct spatial information.

The features extracted by the ResNet and VGG components are concatenated and processed through one GRU layer with 64 units to handle the temporal sequences. The resulting features are then passed through three fully connected (FC) layers with sizes of 1024, 512, and 256, respectively. Three batch normalization and three dropout layers with a rate of 0.5 each follow.

The GRU component is used to model how the FBG signals change over time. It keeps a memory of past measurements and decides which information to keep or forget as new data arrive. This allows the model to learn time-related patterns such as gradual changes in the signal, long-term trends, and consistent growth behavior over time. Although the exact time-related features learned by the GRU cannot be directly observed, its recurrent design allows DIRTNet to capture temporal patterns that cannot be learned using convolutional layers alone.

The output layer of the model was customized for each experiment. In the pseudo-root experiments, two separate models were trained: one to predict root diameter, with an output layer of two neurons representing the two simulated rod diameters, and another to predict root depth, with an output layer of ten neurons representing discrete depth levels.

For the real maize experiments under both control and drought conditions, the model predicted root depth and diameter using an output layer with four neurons representing four output categories. In addition, a separate classification model was trained to distinguish between control and drought conditions using a two-neuron output layer, corresponding to the two classes: control and drought.

### 2.4. Training Methodology

DIRTNet was trained using the same optimization strategy across all experiments. For each experiment, the dataset was split using an 80:20 protocol: 80% of the data was used for training, while the remaining 20% was retained as a hold-out set. The hold-out set was divided equally into validation and test sets. Data augmentation was applied only to the training set to improve model generalization, while the validation and test sets were kept unchanged to ensure fair and unbiased performance evaluation. The validation set was not used for training but was used to monitor loss and F1 scores during training, enabling early stopping and preventing overfitting. Early stopping was applied based on validation loss, with a patience of 10 to 20 epochs. The batch size was fixed at 32, and learning rates were individually optimized within the range of 1 *×* 10*^−^*^6^ to 1 *×* 10*^−^*^3^.

Using the controlled pseudo root experiments, we compared DIRTNet model performance to ten baseline architectures. These baselines included ResNet-18 [30], ResNet-34 [31], VGG-16 [32], VGG-19 [33], Inception-v3 [34], CNN, DenseNet-121 [35], DenseNet-169 [36], LSTM [37], and GRU. This comparison allowed us to identify the strongest competing models under controlled conditions and select the three best-performing baseline models for in-depth evaluation of DIRTNet in the real maize experiments.

## 3. Results

### 3.1. Pseudo-Root Experiments

#### 3.1.1. Data collection and Preprocessing

For generating pseudo-root data, a robotic insertion system was developed to enable controlled penetration of a root-like object into the soil. Two pseudo-roots with diameters of 1 mm and 5 mm were inserted up to a maximum depth of 15 cm below the soil surface over a period of 11 minutes. Ten experimental trials were conducted for each pseudo-root diameter, and optical power signals were continuously recorded from the FBG sensors placed left, right and below the insertion point for dataset construction.

Figure 3 illustrates the FBG power changes induced by the insertion of the 5-mm pseudo-root and the processing steps used to construct the root-depth and root-diameter datasets. The depth dataset was generated from the bottom FBG output, representing depths from 0 cm to 15 cm in 1.5 cm increments. The segment lengths were set to contain a total of 400 data points each, resulting in a total of 300 samples after prepossessing the depth data. The diameter dataset included segments of 300 data points from both side FBGs, resulting in 3,228 samples. To prevent overfitting and improve the model’s generalization capabilities and robustness, we increased the diversity of the training data with augmentation techniques. We opted for adding Gaussian noise [38], scaling [27], shifting [39], flipping [40], and cropping [27, 41] to the segmented signals. After augmentation, the resulting dataset comprised 1,800 depth and 19,368 diameter samples. The augmented dataset was subsequently used to evaluate the performance of our proposed DIRTNet model.

**Figure 3:**
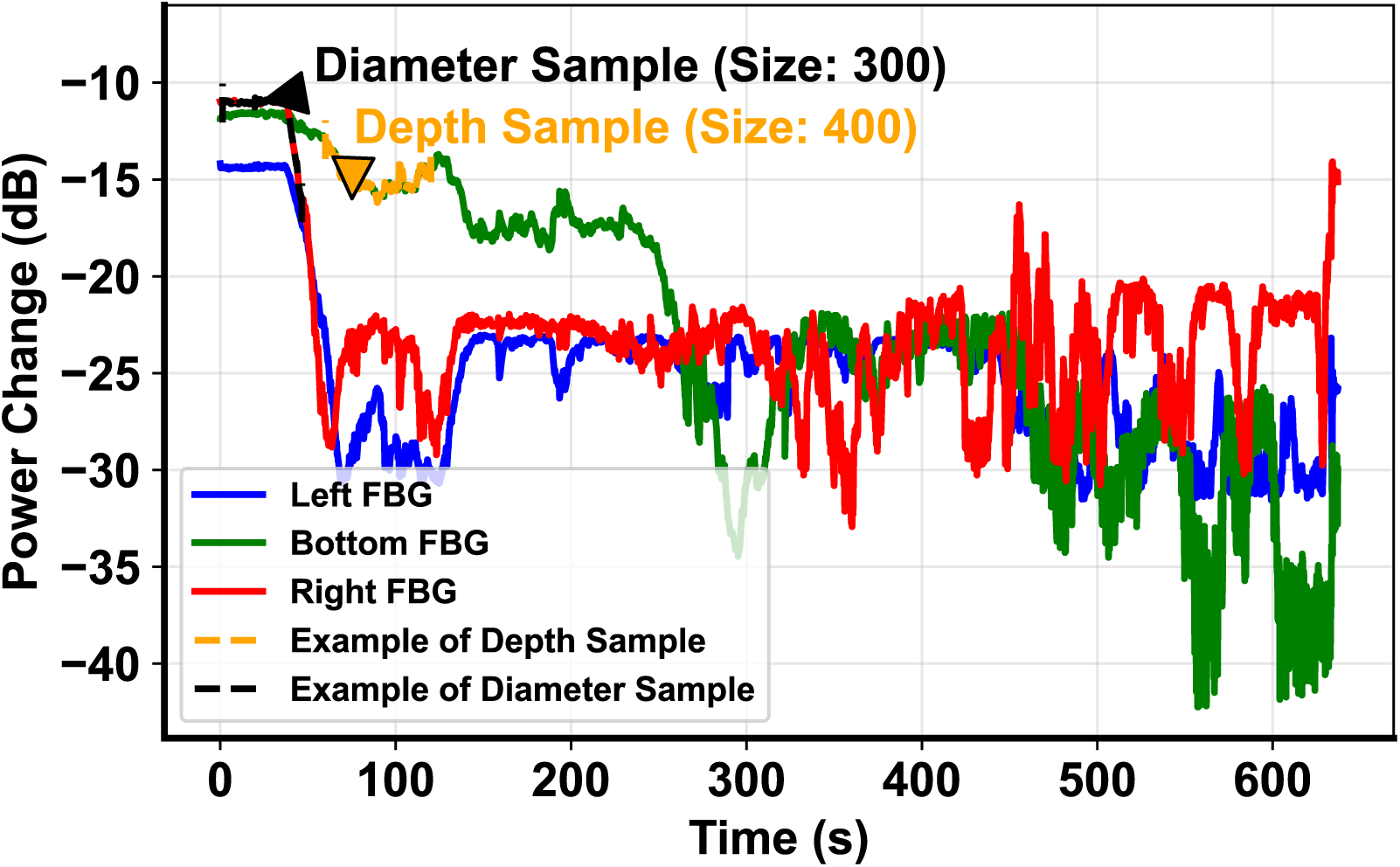
Visualization of FBG Power Changes Induced by 5mm Pseudo-Root, Including Sample Processing Steps for Depth and Diameter Datasets

#### 3.1.2. DIRTNet Evaluation

##### Depth Prediction Model

We determined the training loss to quantify model training efficiency and validation loss to quantify the model generalization potential. As shown in Table 1, the VGG-16, VGG-19, and DenseNet-121 models achieved the lowest validation loss values (0.22, 0.29, and 0.28) combined with low training losses (0.34, 0.56, and 0.32), indicating strong generalization capabilities. In contrast, the ResNet variants had higher training and validation losses, with ResNet-18 having the highest training loss of 0.84 and a high validation loss of 2.95.

**Table 1:**
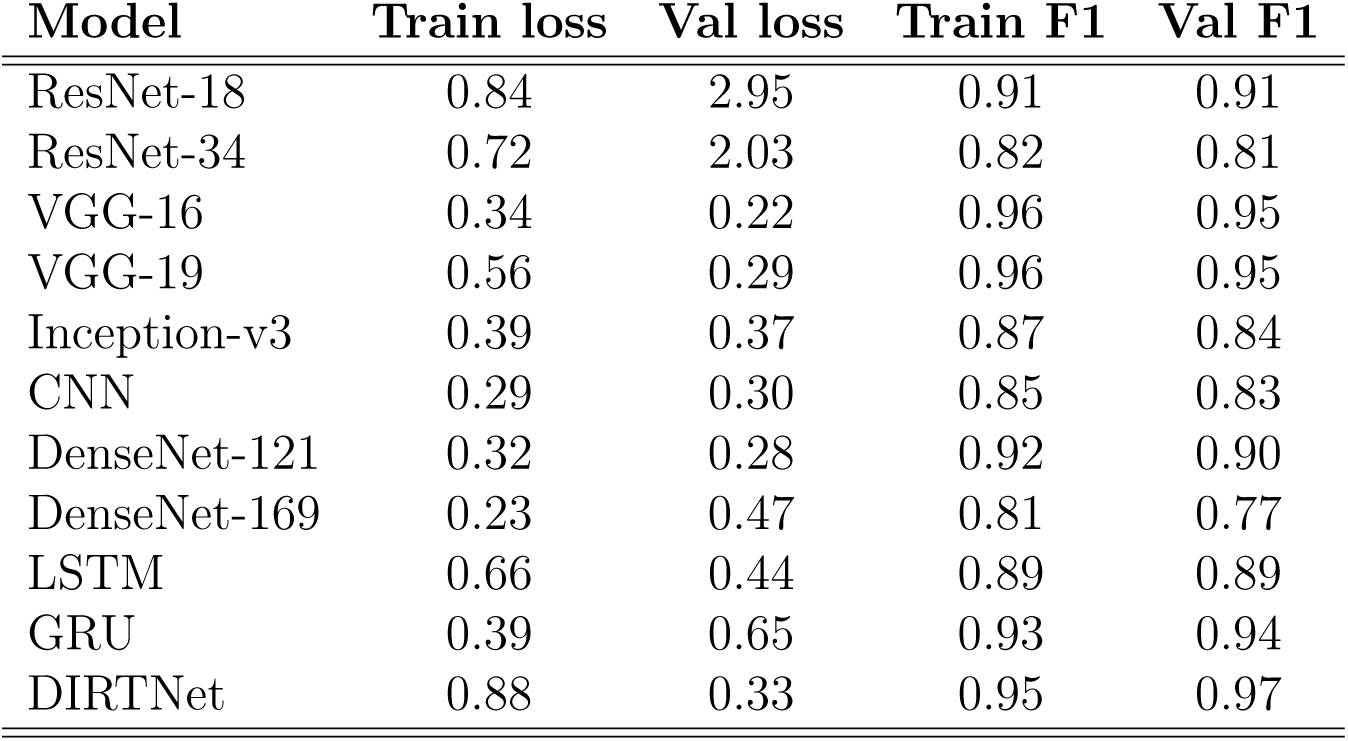
Comparison of Training and Validation Loss, and F1 Scores Across Different Models for Depth Prediction Model.

| **Model** | **Train loss** | **Val loss** | **Train F1** | **Val F1** |
| --- | --- | --- | --- | --- |
| ResNet-18 | 0.84 | 2.95 | 0.91 | 0.91 |
| ResNet-34 | 0.72 | 2.03 | 0.82 | 0.81 |
| VGG-16 | 0.34 | 0.22 | 0.96 | 0.95 |
| VGG-19 | 0.56 | 0.29 | 0.96 | 0.95 |
| Inception-v3 | 0.39 | 0.37 | 0.87 | 0.84 |
| CNN | 0.29 | 0.30 | 0.85 | 0.83 |
| DenseNet-121 | 0.32 | 0.28 | 0.92 | 0.90 |
| DenseNet-169 | 0.23 | 0.47 | 0.81 | 0.77 |
| LSTM | 0.66 | 0.44 | 0.89 | 0.89 |
| GRU | 0.39 | 0.65 | 0.93 | 0.94 |
| DIRTNet | 0.88 | 0.33 | 0.95 | 0.97 |

CNN showed both low validation and training losses (0.30 and 0.29, respectively), indicating effective learning and generalization. Inception-v3 and DenseNet-169 performed moderately, with validation losses of 0.37 and 0.47 and lower training losses of 0.39 and 0.23, respectively. LSTM and GRU also performed moderately, performing similarly in training and validation losses. DenseNet-169, ResNet-34, CNN, Inception-v3, and LSTM showed lower performance, with validation F1 scores ranging from 0.77 to 0.89 and training F1 scores ranging from 0.81 to 0.89.

The proposed hybrid DIRTNet produced a training loss of 0.88 and a substantially lower validation loss of 0.33. This difference likely reflects the use of data augmentation and dropout during training, while the lower validation loss indicates strong generalization to unseen data. To provide a more detailed evaluation of the proposed DIRTNet, we evaluated the training and validation losses for each epoch. Figure 4 (a) shows that during the first few epochs, the training loss decreases smoothly, while the validation loss fluctuates initially. As training progresses from around the 60th to the 100th epoch, both training and validation losses decrease steadily, indicating effective learning. In Figure 4 (b), the training loss decreases smoothly throughout, and the validation loss also decreases, with a gap between training and validation losses emerging at higher epochs. Importantly, this maximum gap remains small, below 0.1–0.2, suggesting good generalization.

**Figure 4:**
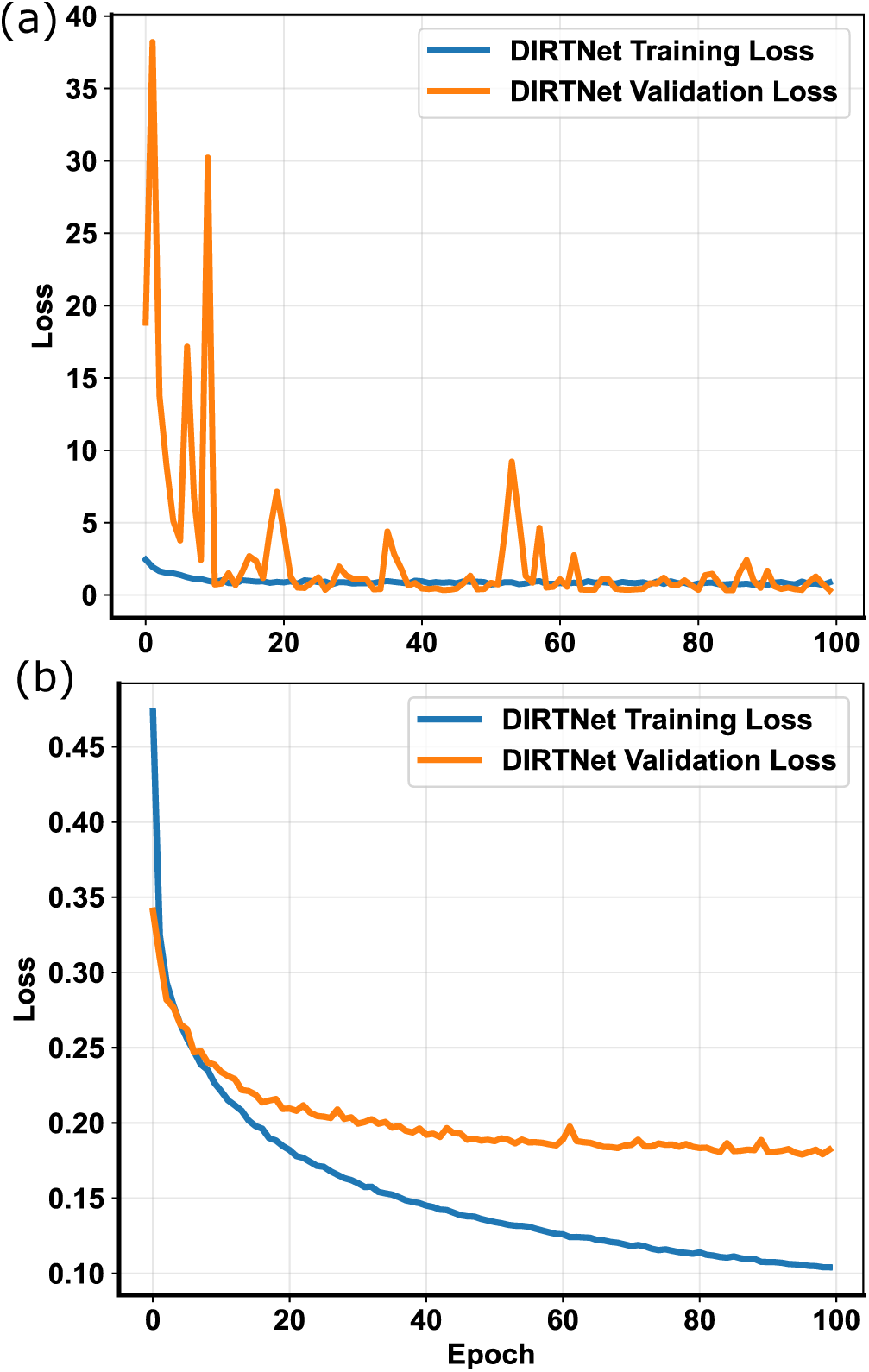
Training and validation loss curves for the proposed DIRTNet. (a) Root depth prediction. (b) Root diameter prediction. These plots illustrate the model’s performance over epochs for both tasks, highlighting trends and convergence behavior.

We also compared F1 scores for training and validation data to quantify model accuracy. As shown in Table 1, VGG-16 and VGG-19 had validation F1 scores of 0.95 each and training F1 scores of 0.96 each. GRU, ResNet-18, and DenseNet-121 achieved moderate F1 scores, ranging from 0.90 to 0.94 for validation and from 0.91 to 0.93 for training. DIRTNet achieved the best validation F1 score of 0.97 and the second-highest training F1 score of 0.95, suggesting excellent performance.

Overall, the evaluation highlights DIRTNet’s superior performance in predicting root depth. It achieves the best balance between training and validation loss, and resulted in the best validation F1 score among the baseline models tested. VGG-16 and VGG-19 also proved to be competitive models for predicting root depth.

##### Diameter Prediction Model

As shown in Table 2, GRU provides the second-best results, with low training and validation losses of 0.18 and 0.23 and F1 scores of 0.94 and 0.92 for the diameter prediction. In contrast, ResNet-34 provides high training and validation losses of 0.73 and 1.80, with F1 scores of 0.60 and 0.80, respectively. The high deviation between the training and validation F1 scores for ResNet-34 indicates potential overfitting. The remaining models, including LSTM, DenseNet-121, DenseNet-169, CNN, Inception-v3, VGG-16, VGG-19, and ResNet-18, show validation losses ranging from 0.29 to 0.44 and training losses ranging from 0.28 to 0.38. Additionally, the training F1 scores range from 0.82 to 0.88, while the validation F1 scores range from 0.80 to 0.88. DIRTNet stands out with the lowest training and validation losses of 0.10 and 0.18 and the highest training and validation F1 scores of 0.97 and 0.94. In Figure 4 (b), the training loss for DIRTNet consistently decreases over epochs, reflecting an improvement in performance as training progresses. The validation loss also shows a steady decline, indicating that the model performs well on unseen data throughout the training process. Although the gap between training and validation losses is higher than current standards, it remains significantly lower than other models tested. DIRTNet is the top-performing model for diameter prediction, demonstrating the lowest training and validation losses and the highest F1 scores. GRU ranks as the second-best model, with competitive performance in both training and validation metrics.

**Table 2:** Comparison of Training and Validation Loss, and F1 Scores Across Different Models for Diameter Prediction Model.

| Model | Train loss | Val loss | Train F1 | Val F1 |
| --- | --- | --- | --- | --- |
| ResNet-18 | 0.37 | 0.44 | 0.82 | 0.84 |
| ResNet-34 | 0.73 | 1.80 | 0.60 | 0.80 |
| VGG-16 | 0.29 | 0.32 | 0.87 | 0.87 |
| VGG-19 | 0.36 | 0.36 | 0.85 | 0.86 |
| Inception-v3 | 0.37 | 0.42 | 0.82 | 0.82 |
| CNN | 0.38 | 0.38 | 0.82 | 0.84 |
| DenseNet-121 | 0.33 | 0.34 | 0.84 | 0.85 |
| DenseNet-169 | 0.33 | 0.32 | 0.85 | 0.80 |
| LSTM | 0.28 | 0.29 | 0.88 | 0.88 |
| GRU | 0.18 | 0.23 | 0.94 | 0.92 |
| DIRTNet | 0.10 | 0.18 | 0.97 | 0.94 |

#### 3.1.3. Performance Metrics Comparison

##### Depth Prediction Model

Table 3 illustrates that the DIRTNet model achieved the highest performance across all metrics, with accuracy, precision, and recall each at 0.97. In the top section of the table, which presents the depth prediction results, DIRTNet clearly outperforms all other models. We also estimated the RMSE for this model to be the lowest at 0.18. This demonstrates its ability to accurately and consistently predict depth, making it the most effective model among those tested. The second-best results were achieved by VGG-16 and VGG-19, which had accuracy, precision, and recall scores of 0.95 each and RMSE values of 0.22.

**Table 3:** Performance metrics (Accuracy, Precision, Recall) of different models for control) and drought c<u>ondition.</u>.

| Model | Accuracy | Precision | Recall |
| --- | --- | --- | --- |
| **Depth** |  |  |  |
| ResNet-18 | 0.91 | 0.92 | 0.91 |
| ResNet-34 | 0.83 | 0.83 | 0.81 |
| VGG-16 | 0.95 | 0.95 | 0.95 |
| VGG-19 | 0.95 | 0.95 | 0.95 |
| DenseNet-121 | 0.90 | 0.90 | 0.90 |
| DenseNet-169 | 0.80 | 0.79 | 0.80 |
| Inception-v3 | 0.87 | 0.86 | 0.86 |
| CNN | 0.83 | 0.84 | 0.85 |
| LSTM | 0.89 | 0.89 | 0.89 |
| GRU | 0.94 | 0.94 | 0.94 |
| DIRTNet | 0.97 | 0.97 | 0.97 |
| **Diameter** |  |  |  |
| ResNet-18 | 0.84 | 0.85 | 0.84 |
| ResNet-34 | 0.80 | 0.82 | 0.80 |
| VGG-16 | 0.87 | 0.87 | 0.87 |
| VGG-19 | 0.86 | 0.86 | 0.86 |
| DenseNet-121 | 0.85 | 0.86 | 0.85 |
| DenseNet-169 | 0.80 | 0.80 | 0.80 |
| Inception-v3 | 0.82 | 0.82 | 0.82 |
| CNN | 0.84 | 0.84 | 0.84 |
| LSTM | 0.88 | 0.88 | 0.88 |
| GRU | 0.92 | 0.92 | 0.92 |
| DIRTNet | 0.94 | 0.94 | 0.94 |

The remaining models, including ResNet-18, ResNet-34, DenseNet-121, DenseNet-169, Inception-v3, CNN, LSTM, and GRU, showed accuracy and recall ranging from 0.80 to 0.94, precision ranging from 0.79 to 0.94, and RMSE ranging from 0.18 to 0.57.

Overall, DIRTNet outperformed all other models in accuracy, precision, recall, and RMSE, demonstrating its superior predictive capability for root depth. VGG-16 and VGG-19 followed as the second-best performers

##### Diameter Prediction Model

Similarly, the bottom part of Table 3, which shows diameter prediction, indicates that DIRTNet outperforms all other models with accuracy, precision, and recall of 0.94 each. Its RMSE was also the lowest at 0.25, demonstrating the model’s strong predictive accuracy and efficiency. The plot also shows that GRU performs notably well, with accuracy, precision, and recall scores of 0.92 each and a relatively low RMSE of 0.29, making it the second-best model. In contrast, ResNet-34 and DenseNet-169 provide lower performance, with accuracy, precision, and recall scores of 0.80 each and the highest RMSE of 0.71. The remaining models, including LSTM, VGG-16, VGG-19, CNN, Inception-v3, and DenseNet-121, achieved accuracy and recall ranging from 0.80 to 0.88, precision ranging from 0.82 to 0.88, and RMSE ranging from 0.34 to 0.44. These results demonstrate DIRTNet’s repeated superior performance in predicting root diameters, with GRU also demonstrating strong performance as the second-best model.

#### 3.1.4. AUC and Confusion Matrix Analysis with DIRTNet

This section presents additional evaluation metrics for DIRTNet, including the Area Under the Curve (AUC) and confusion matrix. The AUC is used for the binary diameter prediction task to measure class separability, while the confusion matrix is used for the multi-class depth prediction task to show detailed class-wise performance. Together, these metrics provide further insight into the model’s performance and reliability.

##### Confusion Matrix for Root Depth Prediction

The confusion matrix provides a detailed breakdown of the actual versus predicted classifications made by the model. The confusion matrix for this task shows the number of correct and incorrect predictions for each class. As shown in Figure 5 (a), a high number of true positives along the diagonal of the matrix indicates that the model is performing well in correctly predicting the root depths. For example, class 0 had 47 correct and 0 incorrect predictions, indicating no prediction errors. For class 8, there were 40 correct and 6 incorrect predictions, with 5 misclassified as class 9 and 1 misclassified as class 7. This indicates that the model performs perfectly for class 0, while for class 8, there is a slight error rate, with some instances being misclassified as classes 7 and 9.

**Figure 5:**
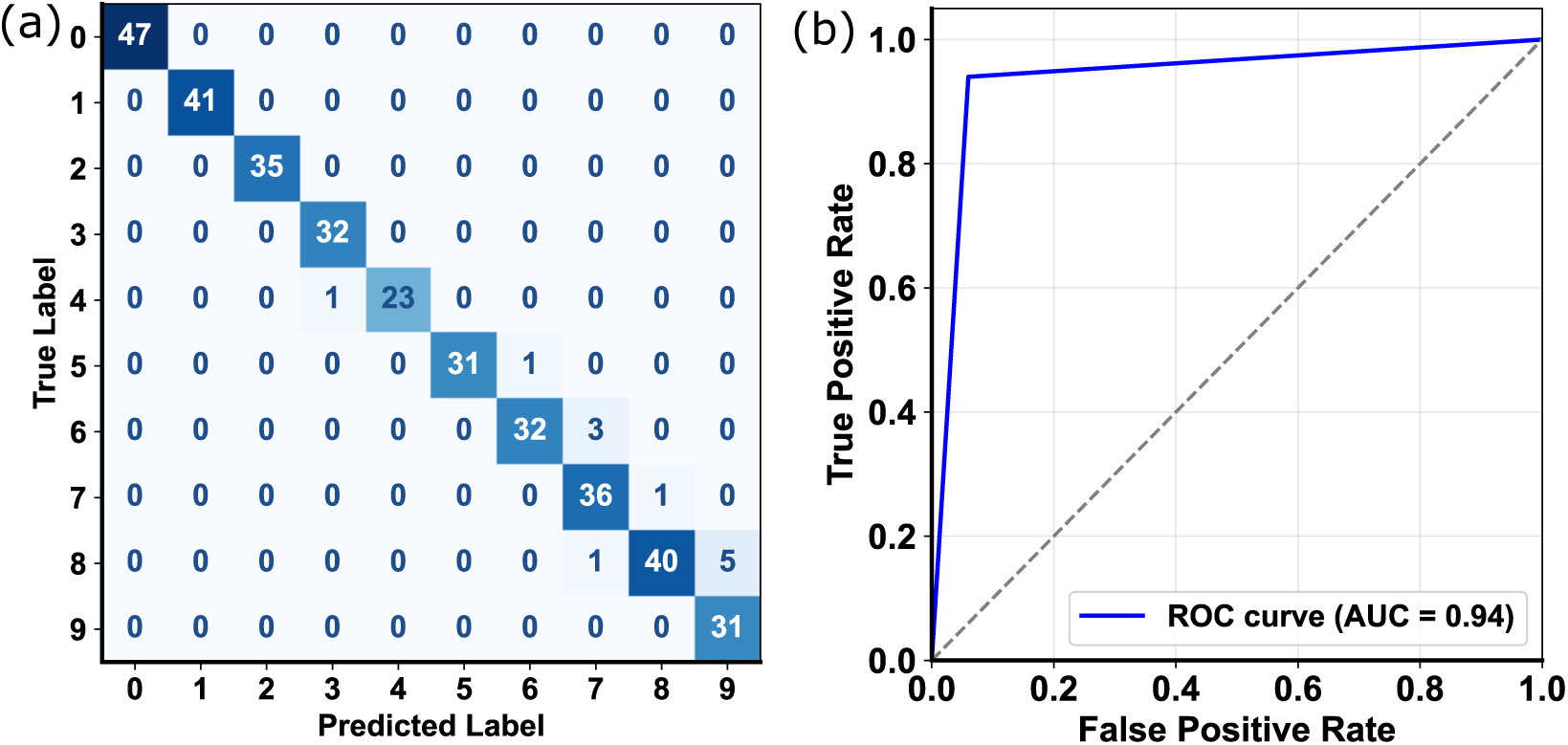
Performance Evaluation of DIRTNet for Root Trait Predictions. (a) Depth Prediction Model(b) Diameter Prediction Model

##### AUC for Root Diameter Prediction

The AUC is a performance measurement for classification models at various threshold settings. A higher AUC value indicates a better performing model. The AUC benefits binary classification tasks by providing a single value that summarizes the performance across all possible classification thresholds. In Figure 5 (b), the proposed DIRTNet achieved an AUC of 0.94. This high AUC value suggests that our model can distinguish between the two classes, indicating excellent performance in predicting root diameters.

### 3.2. Maize Root Experiments

#### 3.2.1. Data collection

We collected FBG sensor data from six sensors embedded in the soil across two potted trials involving a total of 11 maize plants. The first trial consisted of five plants grown entirely under control conditions. The second trial included 6 plants, of which 3 were maintained under well-watered conditions and three were subjected to drought stress. The stress was imposed starting on day 25. For dataset construction, day 1 is considered the first day of observation for all plants.

For Trial 1 (control condition), root depth values up to 10 cm were divided into four equal classes of 2.5 cm each, and root diameter values up to 8 cm were divided into four equal classes of 2 cm each. Plants were monitored from day 1, with variable growth durations across plants. Depth data ranged from day 1 to approximately 34 days for the longest plant and from day 1 to approximately 22 days for the shortest plant. Similarly, diameter data ranged from day 1 to approximately 31 days for the longest plant and from day 1 to approximately 22 days for the shortest plant.

For Trial 2 (drought condition), root depth values up to 20 cm were divided into four equal classes of 5 cm each. Similarly, root diameter values up to 10 cm were divided into four equal classes of 2.5 cm each. Although drought stress was imposed starting on day 25, plants were observed from day 1, and growth durations varied across plants. Depth data ranged from day 1 up to approximately 54 days for the longest plant, and diameter data ranged from day 1 up to approximately 44 days. In both cases, the shortest plants were observed for approximately 34 days. Daily sensor readings were grouped into classes based on the range they fell into, with each class containing multiple daily readings depending on plant growth.

Class ranges for depth and diameter were carefully chosen based on root measurements to optimize model performance across all plants and conditions. The dataset was split using an 80/20 protocol, with 192 initial samples (120 control and 72 drought). Data augmentation was applied only to the training set. This dataset construction approach ensures consistent class labeling across plants and conditions, accounts for variable growth rates, and allows the model to efficiently learn root growth patterns under both control and drought conditions.

#### 3.2.2. Evaluation of Training and Validation Metrics

This analysis focuses on the first experimental trial conducted under control conditions. In the pseudo-root setup, different FBG sensor groups were used for different prediction tasks. The bottom sensor showed stronger sensitivity to vertical strain changes caused by insertion depth and was therefore used for depth prediction, while the left and right sensors captured lateral strain variations more effectively and were used for pseudo-root diameter prediction. These experiments helped identify the spatial sensitivity of different sensor positions and validated the ability of DIRTNet to learn root-related strain patterns from FBG signals. Building on these findings, the real maize experiments used six FBG sensors distributed across upper and lower soil regions to capture more complex biological root-growth behavior. Unlike the pseudo-root setup, where sensor groups were analyzed separately, the maize experiments used signals from all six sensors simultaneously for both root depth and diameter prediction. This multi-sensor design enabled DIRTNet to learn combined spatial and temporal patterns associated with natural root development while maintaining the same architecture and training strategy. Figure 6 shows the training and validation loss curves for DIRTNet. For root depth prediction (Figure 6 (a)), the training loss steadily decreased from 1.34 to 0.57, while the validation loss decreased from 1.35 to approximately 0.65, indicating progressive learning throughout training. For root diameter prediction (Figure 6 (b)), the training loss decreased from 1.39 to 0.13, and the validation loss decreased from 1.39 to approximately 0.15, demonstrating stable convergence. In both cases, the training and validation losses decrease consistently across epochs. Although a small gap between the curves is observed, the validation loss stabilizes toward the later epochs, indicating good generalization without significant overfitting.

**Figure 6:**
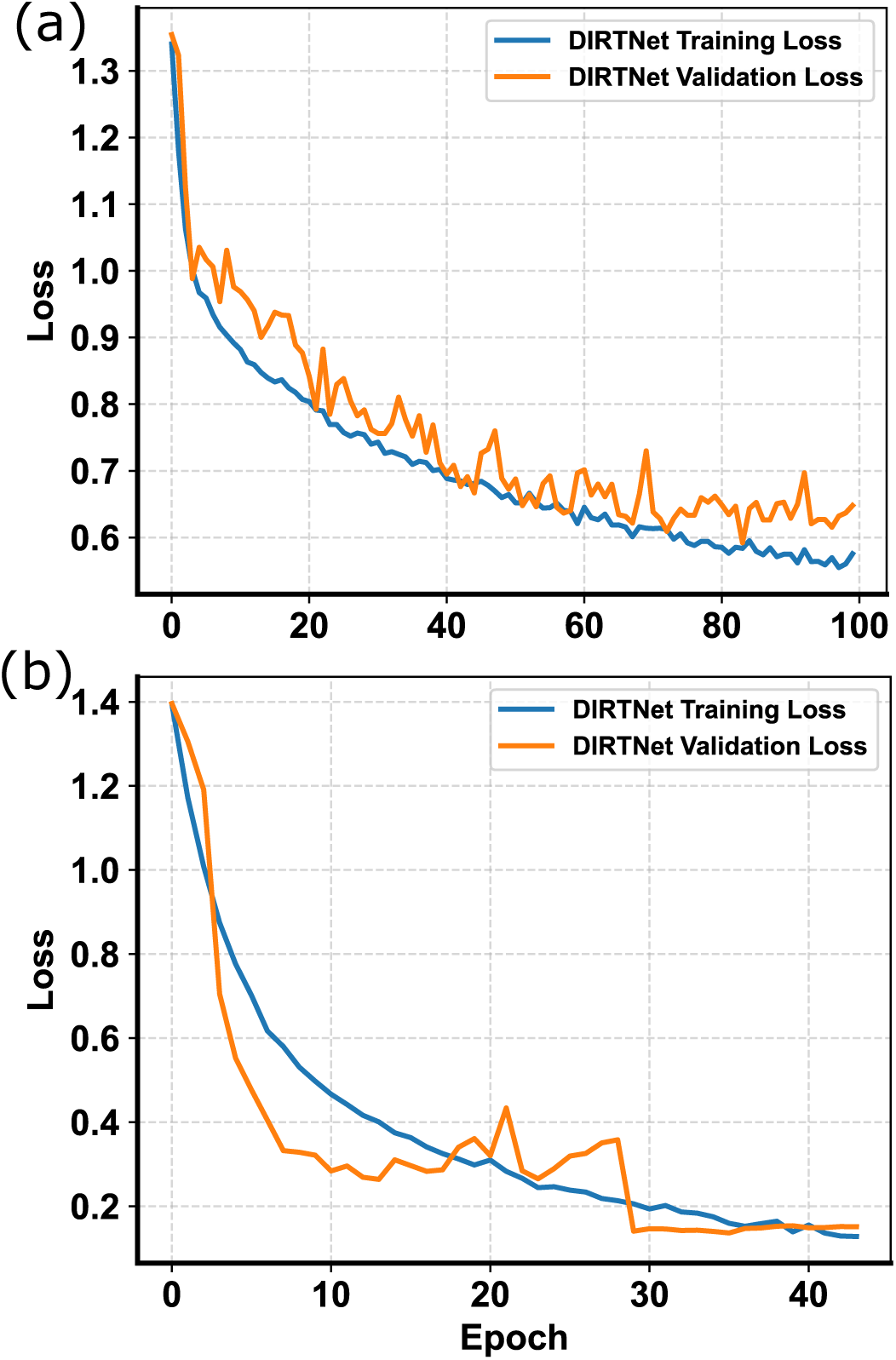
Training and validation loss curves for the proposed DIRTNet using maize root. (a) Root depth prediction. (b) Root diameter prediction. These plots illustrate the model’s performance over epochs for both tasks, highlighting trends and convergence behavior.

It should be noted that training and validation loss curves were not analyzed for the drought experiment, as the drought data were primarily used for evaluating predictive performance and growth-condition classification rather than detailed training-dynamics assessment.

#### 3.2.3. Performance Across Different Growth Durations

To understand how prediction accuracy changes as plants develop over time, we evaluated the DIRTNet model using sensor data from different growth durations under both control and drought conditions. The results are summarized in Figure 7, where Figure 7 (a) shows the control trial and Figure 7 (b) shows the drought trial.

**Figure 7:**
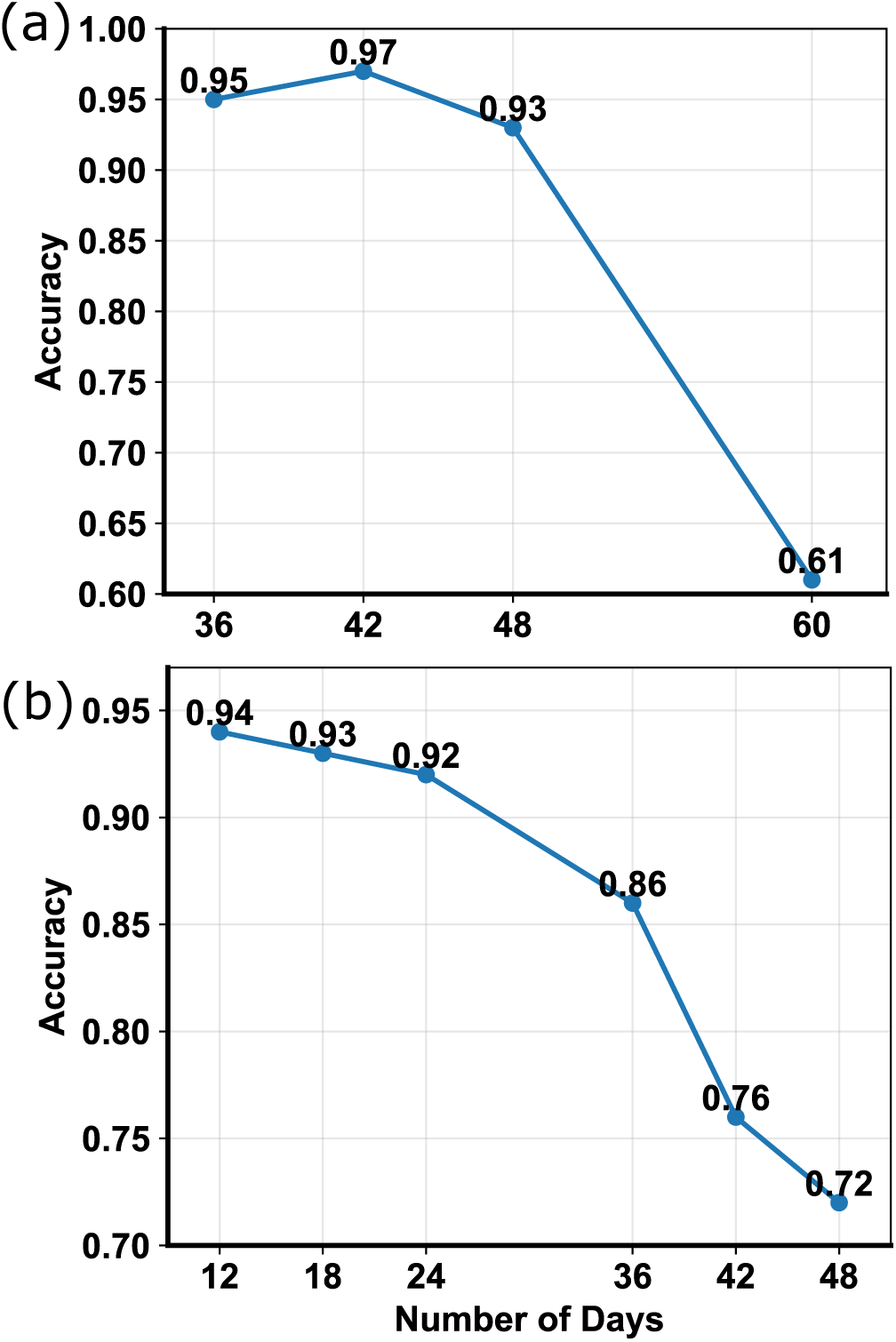
DIRTNet model accuracy across different maize growth stages under (a) control (well-watered) and (b) drought conditions. The model performs best during earlier stages and gradually declines as plant growth and drought stress increase.

For the control trial, data windows of 12, 18, 24, 36, 42, and 48 days were evaluated. As shown in Figure 7 (a), the model achieved the highest accuracy (0.94) using 12 days of data and 0.92 using 24 days. As the observation period increased, the performance gradually declined to 0.86, 0.76, and 0.72 for 36, 42, and 48 days, respectively.

For the drought trial, drought stress was introduced at day 25, and the model was evaluated using data from 36, 42, 48, and 60 days, as shown in Figure 7 (b). The model achieved an accuracy of 0.95 at day 36, which slightly increased to 0.97 at day 42, representing a minor deviation from the general trend. After this point, the performance declined to 0.93 at day 48 and dropped more significantly to 0.61 by day 60. This decrease in accuracy reflects the combined effects of prolonged drought stress and increasing root system complexity, both of which introduce additional variability into the sensor signals and make prediction more difficult.

Overall, these results demonstrate that while DIRTNet performs strongly during earlier growth stages, prediction becomes more challenging as the root structure grows larger and the sensor signals become more complex over time.

#### 3.2.4. DIRTNet Prediction Results

We evaluated the classification performance of the proposed DIRTNet model for both root depth and diameter under control and drought conditions.

##### Control Conditions

The DIRTNet model demonstrated strong performance in predicting both root depth and diameter, as shown in Figure 8. Figure 8 (a) presents the comparison of predicted and actual depth values, while Figure 8 (b) shows the comparison of predicted and actual diameter values. Overall, the model achieved an accuracy of 0.92 for both traits, with precision and recall values of 0.94 and 0.92, respectively. The predicted values closely follow the actual class distributions, indicating that the model reliably distinguishes between the different classes for both depth and diameter. Minor discrepancies are observed at certain boundary classes, but these are limited and do not significantly affect overall performance. These results suggest that the model can effectively capture root width and depth from the input data, supporting its potential use for automated root phenotyping. **Drought Conditions:** In the drought experiment, the DIRTNet model demonstrated strong performance in predicting both root depth and diameter, achieving an accuracy of 0.92 with precision and recall of 0.94 and 0.92, respectively. Although the actual versus predicted curves are not shown, the model exhibited trends similar to those observed under control conditions, indicating consistent and reliable prediction of root traits under drought stress. These results support the potential of DIRTNet for automated phenotyping across diverse environmental conditions.

**Figure 8:**
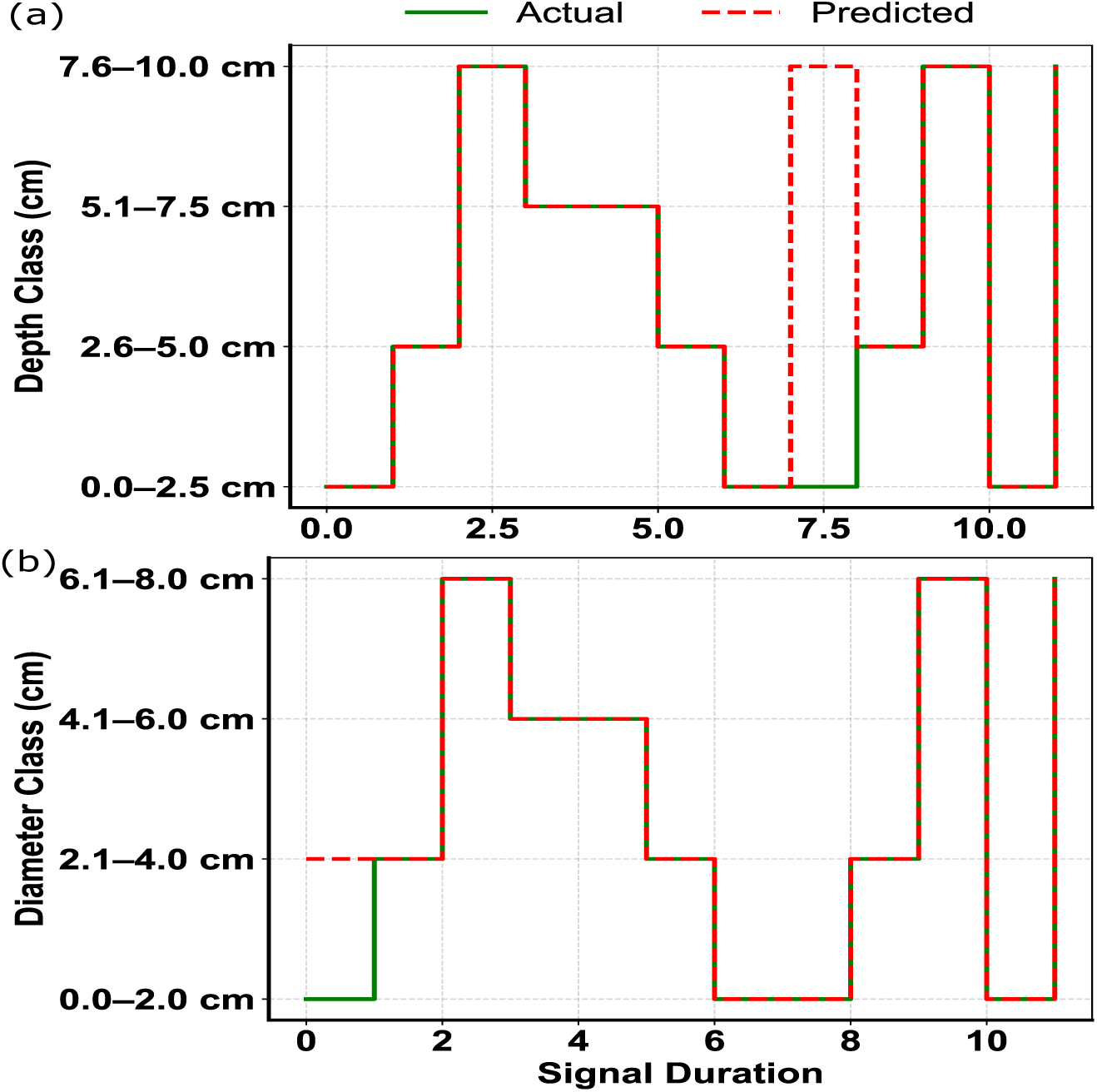
Step plot comparison of DIRTNet predictions vs. actual measurements for root growth: (a) depth prediction (0–10 cm, 4 classes), (b) diameter prediction (0–8 cm, 4 classes).

In general, we observed that as the number of days in the dataset increases, the data become more complex, making prediction more challenging. This behavior is likely caused by continued root growth around the sensor, which weakens the signal, making it less informative for prediction. However, in the drought experiment, even though more days of data were included, the model achieved performance similar to the control. One reason could be that for each class, the longer length intervals in drought corresponded to a larger number of days per class, providing the model with more information for each prediction. In contrast, in the control data, each class covered fewer days, giving less information and making predictions slightly more difficult.

#### 3.2.5. Performance Metrics Comparison

We compared ResNet-18, VGG-16, and DIRTNet using the control maize dataset for predicting root depth and diameter. As summarized in Table 4, DIRTNet outperformed both ResNet-18 and VGG-16, achieving the highest accuracy and recall of 0.92, with a precision of 0.94 for both traits. ResNet-18 showed moderate performance across all metrics for depth and diameter prediction, while VGG-16 exhibited moderate performance for diameter but poor performance for depth. Recurrent-only models such as LSTM and GRU performed poorly on their own. However, incorporating GRU within the DIRTNet architecture, together with spatial feature extraction, enabled the model to capture both temporal and spatial patterns, resulting in strong overall performance.

**Table 4:** Comparison of different models for Depth and Diameter prediction under control conditions in terms of Accuracy, Precision, and Recall.

| Model | Accuracy | Precision | Recall |
| --- | --- | --- | --- |
| **Depth** |  |  |  |
| ResNet18 | 0.83 | 0.92 | 0.83 |
| VGG16 | 0.67 | 0.73 | 0.67 |
| DIRTNet | 0.92 | 0.94 | 0.92 |
| **Diameter** |  |  |  |
| ResNet18 | 0.83 | 0.90 | 0.83 |
| VGG16 | 0.83 | 0.90 | 0.83 |
| DIRTNet | 0.92 | 0.94 | 0.92 |

Since DIRTNet consistently outperformed all other models on the control dataset, it was selected as the best-performing model and evaluated on the drought data. The other baseline models were not tested on drought conditions, as DIRTNet had already been established as the top-performing architecture. This approach allows us to focus on evaluating DIRTNet’s ability to predict root traits under early drought stress without redundant testing of weaker models.

### 3.3. DIRTNet Classification Results for Control and Drought Conditions

We tested DIRTNet’s ability to classify plant growth conditions (nonlimiting vs. drought) using FBG sensor data from two different setups. In both setups, the first 24 days of data were considered non-limiting conditions, and drought conditions were induced from day 25, with drought data collected up to day 48.

In the first setup, data from eleven plants were used, eight from the non-limiting trial and three from the drought trial. After augmentation, the training set contained 21,726 samples. We achieved a test accuracy and recall of 0.87, with precision of 0.84. In the second setup, data from three plants from the second trial (drought only) were used. The model achieved a test accuracy of 0.88, with precision of 0.86 and recall of 0.90.

We found that DIRTNet can reliably classify non-limiting and drought conditions based on RSA using in-soil FBG signals. The model performed well across both environments, showing that it effectively captures stress-induced changes in root growth in or experiments.

## 4. Discussion

Our team previously characterized FBG fiber optic sensors and their potential for monitoring underground root morphology (such as width and depth) and environmental factors such as groundwater levels [24]. The studies highlighted the sensor’s non-destructive, continuous monitoring capabilities and potential agricultural and environmental applications. Our earlier work [23] utilized a ResNet-based deep learning network to evaluate the sensor output for non-destructive root monitoring, which revealed limitations of ResNet-based approaches because of the combined spatio-temporal signal analysis needed for root phenotyping applications. The presented work overcomes the observed limitations by introducing DIRTNet, a hybrid Deeplearning model designed for plant root phenotyping using FBG sensor data.

In the simulation experiments, we used metal rods of different diameters to generate controlled pseudo-root signals. This setup allowed us to systematically benchmark DIRTNet against ten baseline models. Across all metrics, DIRTNet outperformed the baseline models for predicting root depth and diameter. For depth prediction, DIRTNet achieved balanced training and validation losses of 0.88 and 0.33, with F1 scores of 0.95 and 0.97, indicating strong and consistent performance. Among ResNet variants, ResNet-18 had the highest training and validation losses (0.84 and 2.95) but still achieved higher F1 scores than ResNet-34. This can be explained by the possibility that ResNet-34 might not be fitting well, struggling to capture the most relevant patterns in the data. Despite its relatively lower loss, it may not be capturing the data as effectively as ResNet-18, leading to lower F1 scores. For diameter prediction, DIRTNet achieved 0.10 training and 0.18 validation losses with 0.94 accuracy, precision, and recall, and an RMSE of 0.25, showing balanced performance. By contrast, ResNet-34 had higher losses (0.73 training, 1.80 validation) and a larger discrepancy between training and validation F1 scores, indicating less stable performance.

For real maize plants under control conditions, we compared the top-performing models from simulation (DIRTNet, VGG, and ResNet) using data from first trial (five control plants). DIRTNet achieved the highest performance for both depth and diameter, with accuracy and recall of 0.92 and precision of 0.94 for both traits.

We also evaluated recurrent-only models, LSTM and GRU, in both the simulation and maize root experiments. In the simulation study, where the two rods grew uniformly and consistently in length and downward direction, LSTM and GRU performed well due to strong and predictable temporal patterns. However, in real maize roots, recurrent-only models performed poorly when used alone. This is likely because individual plants grow heterogeneously, some roots grow faster or taller, while others remain shorter, resulting in inconsistent temporal patterns that recurrent-only models cannot reliably capture. In DIRTNet, GRU is combined with spatial feature extraction, allowing the model to exploit temporal patterns effectively only when paired with spatial context, resulting in robust and accurate root trait prediction under heterogeneous growth conditions.

Training on early growth data provided the best results, while performance declined gradually with longer observation windows (e.g., 0.72 accuracy by day 48) due to increasing signal complexity. The proposed models, improved efficiency and generalization. For drought conditions, DIRTNet achieved peak accuracy of 0.97 at Day 42, with a slight drop to 0.93 by Day 48. This gradual decrease in accuracy as the days progress may be attributed to increasing complexity in root growth patterns, overlapping root signals, and higher environmental variability that introduce noise into the FBG sensor measurements. Additionally, as roots grow larger and interact more dynamically with the surrounding soil, the sensor signals become more challenging to interpret, which can slightly reduce model performance. One potential solution to mitigate this issue is to extend the training dataset to include later-stage growth signals.

For control vs. drought classification, using combined FBG signals from both trials, DIRTNet was again the only model tested. On 11 plants (8 control + 3 drought), it achieved 0.87 accuracy, 0.84 precision, and 0.87 recall. Using drought-only data from three plants, accuracy improved to 0.88, with precision 0.86 and recall 0.90. This demonstrates that DIRTNet can reliably distinguish growth conditions based on in-soil root signals.

Overall, DIRTNet consistently showed strong predictive performance across simulation, control, and drought experiments. Its hybrid design allows it to handle complex, time-dependent FBG signals, accurately predicting root traits and classifying stress conditions. Limitations include reduced performance for longer growth periods, and a relatively small number of plants, which may affect generalizability. However, our study was limited to distributing sensors in the 5 gallon pots and maize roots spread further than the pot diameter. While we can not limit the growth of the root, we could limit the data used to a range where the root did not outgrew the distance to the sensor.

Despite these limitations, DIRTNet is a key foundation advancing fiber optics sensing towards continuous, non-destructive in situ root phenotyping to support the development of climate-resilient crops.

## 5. Conclusion and Future Work

A non-destructive approach for root phenotyping using Fiber Bragg Grating (FBG) sensors combined with neural networks is presented, providing an effective way to quantify root growth and development under both control and drought conditions. We developed DIRTNet, a hybrid deep learning model that combines ResNet and VGG for spatial feature extraction with GRU for temporal sequence processing, enabling highly accurate predictions of root diameter and depth.

For drought conditions, only DIRTNet was evaluated, because it was the top-performing model in control experiments. Using drought data, DIRTNet achieved high accuracy, precision, and recall in predicting root traits, demonstrating its ability to capture stress-induced changes in root development. Furthermore, DIRTNet successfully classified control versus drought conditions, confirming its effectiveness in monitoring root responses to environmental stress.

These results confirm that the hybrid architecture effectively combines spatial and temporal information from FBG sensors, making DIRTNet a reliable tool for non-destructive root phenotyping. Future work will focus on optimized 3D sensor distribution in the soil to enable growth periods over the whole life cycle of the plant, extending the model to agricultural production fields that include tractors and other machinery influencing the signals. We envision that further integration with other phenotyping and sensing platforms will enable large-scale root phenotyping of entire fields throughout plant development. Therefore, DIRTNet is a critical component to support climate-resistant crop development and improved phenome characterization and phene discovery [42].

## Acknowledgment

This work was supported by the National Science Foundation under CAREER Award 2329282.

## References

[1] U. Nations,Desertification and drought day 2024: “united for land: Our legacy. our future”, 2024, accessed: (18. september 2026).

[2] R. G. Araújo, R. A. Chavez-Santoscoy, R. Parra-Saldívar, E. M. Melchor-Martínez, H. M. Iqbal, Agro-food systems and environment: Sustaining the unsustainable, Current Opinion in Environmental Science & Health 31 (2023) 100413.

[3] J. P. Lynch, Edaphic stress interactions: important yet poorly understood drivers of plant production in future climates, Field Crops Research 283 (2022) 108547.

[4] P. Singh, A. Sharma, J. Dhankhar, Climate Change and Soil Fertility, Springer Nature Singapore, Singapore, 2022, pp. 25–59. doi:10.1007/978-981-16-7759-5_3_. *URL*

[5] R. Lal, Climate change and soil degradation mitigation by sustainable management of soils and other natural resources, Agric Res 1 (2012) 199–212. doi:10.1007/s40003-012-0031-9.

[6] B. Gruber, R. Giehl, S. Friedel, N. von Wirén, Plasticity of the arabidopsis root system under nutrient deficiencies, Plant Physiology 163 (2013) 161–179.

[7] K. Yamazaki, T. Fujiwara, The effect of phosphate on the activity and sensitivity of nutritropism toward ammonium in rice roots, Plants 11 (6) (2022).

[8] M. Mickelbart, P. Hasegawa, J. Bailey-Serres, Genetic mechanisms of abiotic stress tolerance that translate to crop yield stability, Nature Reviews Genetics 16 (4) (2015) 237–251.

[9] A. Bucksch, A. Atta-Boateng, A. F. Azihou, D. Battogtokh, A. Baumgartner, B. M. Binder, S. A. Braybrook, C. Chang, V. Coneva, T. J. DeWitt, et al., Morphological plant modeling: unleashing geometric and topological potential within the plant sciences, Frontiers in plant science 8 (2017) 900.

[10] R. Metzner, A. Eggert, D. van Dusschoten et al., Direct comparison of mri and x-ray ct technologies for 3d imaging of root systems in soil: potential and challenges for root trait quantification, Plant Methods 11 (2015). doi:10.1186/s13007-015-0060-z. URL https://doi.org/10.1186/s13007-015-0060-z

[11] R. Flavel, C. Guppy, M. T., et al., Non-destructive quantification of cereal roots in soil using high-resolution x-ray tomography, Journal of Experimental Botany 63 (2012) 2503–2511.

[12] R. Flavel, C. Guppy, S. Rabbi, I. Young, An image processing and analysis tool for identifying and analyzing complex plant root systems in 3-d soil using non-destructive analysis: Root1, PLoS ONE 10 (2017) 1–18.

[13] J. Lafond, L. Han, P. Duiteul, Concepts and analyses in the ct scanning of root systems and leaf canopies: a timely summary, Frontiers in Plant Science 6 (2015) 1–7.

[14] X. Liu, X. Dong, D. Leskovar, Ground penetrating radar for underground sensing in agriculture: a review, International Agrophysics 30 (4) (2016) 533–543.

[15] J. Burridge, et al., Legume shovelomics: High-throughput phenotyping of common bean (phaseolus vulgaris) and cowpea (vigna unguiculata) root architecture in the field, Field Crops Research 192 (2016) 21–32.

[16] J. Kengkanna, et al., Phenotypic variation of cassava root traits and their responses to drought, Applications in Plant Sciences 7 (2019).

[17] A. Bucksch, J. Burridge, L. M. York, A. Das, E. Nord, J. S. Weitz, J. P. Lynch, Image-based high-throughput field phenotyping of crop roots, Plant Physiology 166 (2) (2014) 470–486.

[18] S. Liu, C. S. Barrow, M. Hanlon, J. P. Lynch, A. Bucksch, Dirt/3d: 3d root phenotyping for field-grown maize (zea mays), Plant physiology 187 (2) (2021) 739–757.

[19] R. Soman, J. Wee, K. Peters, Optical fiber sensors for ultrasonic structural health monitoring: A review, Sensors 21 (21) (2021). doi:10.3390/s21217345. URL https://www.mdpi.com/1424-8220/21/21/7345

[20] H. Guo, G. Xiao, N. Mrad, J. Yao, Fiber optic sensors for structural health monitoring of air platforms, Sensors 11 (4) (2011) 3687–3705. doi:10.3390/s110403687. URL https://www.mdpi.com/1424-8220/11/4/3687

[21] M. Tei, F. Soma, E. B., et al., Non-destructive real-time monitoring of underground root development with distributed fiber optic sensing, Plant Methods 20 (2024). doi:10.1186/s13007-024-01160-z. URL https://doi.org/10.1186/s13007-024-01160-z

[22] S. Binder, K. Hossain, A. Bucksch, M. Fok, Non-destructive underground fiber Bragg grating sensing system with ResNet prediction for root phenotyping, in: F. Ferranti, M. K. Hedayati, A. Fratalocchi (Eds.), Machine Learning in Photonics, Vol. 13017, International Society for Optics and Photonics, SPIE, 2024, p. 1301703. doi:10.1117/12.3016810. URL https://doi.org/10.1117/12.3016810

[23] S. Binder, K. Hossain, A. Bucksch, M. Fok, Fiber bragg grating-based sensing system for non-destructive root phenotyping with resnet prediction, IEEE Photonics Technology Letters 37 (8) (2025) 473–476. doi:10.1109/LPT.2025.3553701.

[24] S. Binder, M. Yang, Q. Victor, et al., Non-destructive measurements of root traits and their soil-water environment using fiber bragg grating-based fiber optic sensors, ESS Open Archive (November 2021). URL https://essopenarchive.org/

[25] A. Krizhevsky, I. Sutskever, G. Hinton, Imagenet classification with deep convolutional neural networks, in: Advances in Neural Information Processing Systems, 2012, pp. 1097–1105.

[26] K. Simonyan, A. Zisserman, Very deep convolutional networks for large-scale image recognition, CoRR (2014).

[27] K. He, X. Zhang, S. Ren, J. Sun, Deep residual learning for image recognition, in: 2016 IEEE Conference on Computer Vision and Pattern Recognition (CVPR), IEEE, Las Vegas, NV, USA, 2016, pp. 770–778.

[28] J. Chung, C. Gulcehre, K. Cho, Y. Bengio, Empirical evaluation of gated recurrent neural networks on sequence modeling, arXiv preprint (2014).

[29] G. Meltz, K. Hill, Fiber bragg grating technology fundamentals and overview, Journal of Lightwave Technology 15 (1997) 1263 – 1276.

[30] K. He, X. Zhang, S. Ren, J. Sun, Deep residual learning for image recognition, in: Proceedings of the 2016 IEEE Conference on Computer Vision and Pattern Recognition (CVPR), 2016, pp. 770–778. doi:10.1109/CVPR.2016.90.

[31] A. A. Rizaldi, E. Gautama, L. Kamelia, Performance measurement of resnet-34 in convolutional neural network method for classification of mask type usage, in: Proceedings of the 2023 6th International Conference of Computer and Informatics Engineering (IC2IE), 2023, pp. 115–120. doi:10.1109/IC2IE60547.2023.10331025.

[32] J. Ha, H. Moon, J. Kwak, S. Hassan, L. Dang, O. Lee, H. Park, Deep convolutional neural network for classifying fusarium wilt of radish from unmanned aerial vehicles, Journal of Applied Remote Sensing 11 (2017) 042621.

[33] L. A. Gatys, A. S. Ecker, M. Bethge, Image style transfer using convolutional neural networks, in: Proceedings of the IEEE Conference on Computer Vision and Pattern Recognition (CVPR), 2016, pp. 2414– 2423.

[34] L. Dang, S. Hassan, I. Suhyeon, A. Sangaiah, I. Mehmood, S. Rho, S. Seo, H. Moon, I. Syed, Uav based wilt detection system via convolutional neural networks, Sustainable Computing: Informatics and Systems (2018).

[35] S. A. Albelwi, Deep architecture based on densenet-121 model for weather image recognition, International Journal of Advanced Computer Science and Applications 13 (10) (2022). doi:10.14569/IJACSA.2022.0131065. URL https://doi.org/10.14569/IJACSA.2022.0131065

[36] R. Joshi, P. Negi, T. Poongodi, Multilabel classifier using densenet-169 for alzheimer’s disease, in: 2023 4th International Conference on Intelligent Engineering and Management (ICIEM), 2023, pp. 1–7. doi:10.1109/ICIEM59379.2023.10165844.

[37] S. Hochreiter, J. Schmidhuber, Long short-term memory, Neural Computation 9 (8) (1997) 1735–1780.

[38] M. Sáiz-Abajo, B.-H. Mevik, V. Segtnan, T. Næs, Ensemble methods and data augmentation by noise addition applied to the analysis of spectroscopic data, Analytica Chimica Acta 533 (2) (2005) 147–159. 10.1016/j.aca.2004.10.086.

[39] M. B. Er, I. B. Aydilek, Music emotion recognition by using chroma spectrogram and deep visual features, International Journal of Computational Intelligence Systems 12 (2) (2019) 1622–1634. doi:10.2991/ijcis.d.191216.001.

[40] L. Huang, W. Pan, Y. Zhang, L. Qian, N. Gao, Y. Wu, Data augmentation for deep learning-based radio modulation classification, IEEE Access 8 (2019) 1498–1506.

[41] C. Szegedy, W. Liu, Y. Jia, P. Sermanet, S. Reed, D. Anguelov, D. Erhan, V. Vanhoucke, A. Rabinovich, Going deeper with convolutions, in: Proceedings of the IEEE Conference on Computer Vision and Pattern Recognition (CVPR), IEEE, 2015, pp. 1–9.

[42] A. Bucksch, Y. S. Chung, J. L. Clarke, S. Gerth, P. von Gillhaussen, W. Guo, J. Kholová, S. Pariyar, E. Pickering, S. Sankaran, et al., Plant phenomics—the unrecognized rise of a scientific discipline, Trends in Plant Science (2026).

